# A theory of rapid behavioral inferences under the pressure of time

**DOI:** 10.1101/2024.08.26.609738

**Authors:** Ann M. Hermundstad, Wiktor F. Młynarski

## Abstract

To survive, animals must be able quickly infer the state of their surroundings. For example, to successfully escape an approaching predator, prey must quickly estimate the direction of approach from incoming sensory stimuli and guide their behavior accordingly. Such rapid inferences are particularly challenging because the animal has only a brief window of time to gather sensory stimuli, and yet the accuracy of inference is critical for survival. Due to evolutionary pressures, nervous systems have likely evolved effective computational strategies that enable accurate inferences under strong time limitations. Traditionally, the relationship between the speed and accuracy of inference has been described by the “speed-accuracy tradeoff” (SAT), which quantifies how the average performance of an ideal observer improves as the observer has more time to collect incoming stimuli. While this trial-averaged description can reasonably account for individual inferences made over long timescales, it does not capture individual inferences on short timescales, when trial-to-trial variability gives rise to diverse patterns of error dynamics. We show that an ideal observer can exploit this single-trial structure by adaptively tracking the dynamics of its belief about the state of the environment, which enables it to speed its own inferences and more reliably track its own error, but also causes it to violate the SAT. We show that these features can be used to improve overall performance during rapid escape. The resulting behavior qualitatively reproduces features of escape behavior in the fruit fly *Drosophila melanogaster*, whose escapes have presumably been highly optimized by natural selection.

## INTRODUCTION

An animal’s survival depends on its ability to successfully interact with its surroundings. This requires continually inferring properties of the environment from incoming sensory stimuli to guide appropriate actions. Inference is thus a fundamental computation that must be performed by the nervous system, often in challenging circumstances that are limited by noise [1] and metabolic [2] or computational [3, 4] constraints.

Of the many constraints that make inference challenging, strong time limitations are perhaps the most directly related to animal’s survival. For example, prey must be able to estimate the direction of an approaching predator and plan an effective escape strategy, often in a fraction of a second [5, 6]. In such scenarios, accurate inference is challenging because the brain has limited time to gather sensory data that can itself be noisy and inaccurate. Because of the direct impact of rapid inference on life and death decisions, brains and other biological systems have likely evolved sophisticated strategies to maximize inference accuracy when faced with strong time limitations [7, 8].

Canonical theoretical approaches use the speed-accuracy-tradeoff (SAT) to describe the behavior of an ideal observer that must balance the time spent collecting sensory stimuli with the accuracy of the resulting inference [9, 10]. The SAT has an intuitive interpretation: the longer the observer waits, the more sensory stimuli it can gather, and the more accurate the inference it can make. To determine when to stop collecting sensory stimuli and commit to an action, the observer must thus trade off the speed of its decision against the accuracy of its inference. This tradeoff has been shown to capture behavior across multiple scenarios and species, including in humans [11–14], rats [15], mice [16], bees [17], and non-neural organisms such as the slime mold *Physarum polycephallum* [18].

The SAT establishes a fundamental relationship between the speed and accuracy of inference, and suggests that animals should use all available time to achieve the highest possible accuracy. However, the SAT characterizes the performance of an ideal observer *on average*, and it ignores any structure observed in individual scenarios or trials. In particular, short sequences of stimuli can result in either much better or much worse performance than is captured by the average. In principle, this trial-to-trial structure could be exploited by detecting stimulus sequences that lead to a rapid reduction in inference error, thereby enabling the observer to reduce inference time and devote more time to coordinating actions. Alternatively, this structure could be exploited by detecting stimulus sequences that are likely to result in high inference errors that lead to inaccurate beliefs, thereby enabling the observer to take actions that maximize the chance of survival even when the underlying state of the environment is unknown (e.g. as observed in freezing behavior [6]). In both cases, the ability to exploit higher-order statistics of individual inferences could give animals a richer repertoire of possible actions and paths to survival, beyond waiting for more data or acting with inaccurate estimates, as suggested by the SAT.

In this work, we propose a theory for how an ideal observer can exploit the structure of trial-to-trial variability in order to increase the speed of its inference and bound the expected magnitude of its error. Our key insight is that short sequences of stimuli generate diverse patterns of error on individual trials; the observer can exploit this structure by relying on the dynamics of its own uncertainty about the underlying state of the environment. As a result, the optimal adaptive observer violates the SAT. Using a simple observer model that mimics an escape task, we show that this adaptive inference strategy improves overall escape performance, and qualitatively captures features of escape behavior in *Drosophila melanogaster*. This suggests that these and other organisms might exploit statistical regularities to improve inferences under strong time constraints.

## RESULTS

Our primary conceptual aim is to characterize trial-to-trial variability that arises when performing inference under strong time constraints, and identify statistical structure that could be exploited by an ideal observer to improve the speed and reliability of inference. We first demonstrate the existence of structured variability in a simple estimation task: inferring the mean of a Gaussian distribution from a limited number of samples. We then demonstrate how animals could exploit such variability to improve their performance in a time-constrained task that is crucial for survival: rapid escape.

### Inference tasks exhibit structured variability in their trial-to-trial dynamics

In an inference task, an organism estimates a latent state of the environment *θ* from a sequence of incoming sensory stimuli *s*_1_, …, *s*_*t*_ (Fig. 1a). The organism can be modeled as an ideal observer that performs optimal inference by computing the posterior distribution over the latent state, given the observed stimulus sequence (i.e., by computing *p*(*θ*|*s*_*τ*≤*t*_), where *s*_*τ*≤*t*_ is the specific sequence of stimuli observed in a trial up to time point *t*). This posterior can then be used to compute a point estimate of the latent state 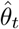, for example by taking the posterior average [19].

**Figure 1:**
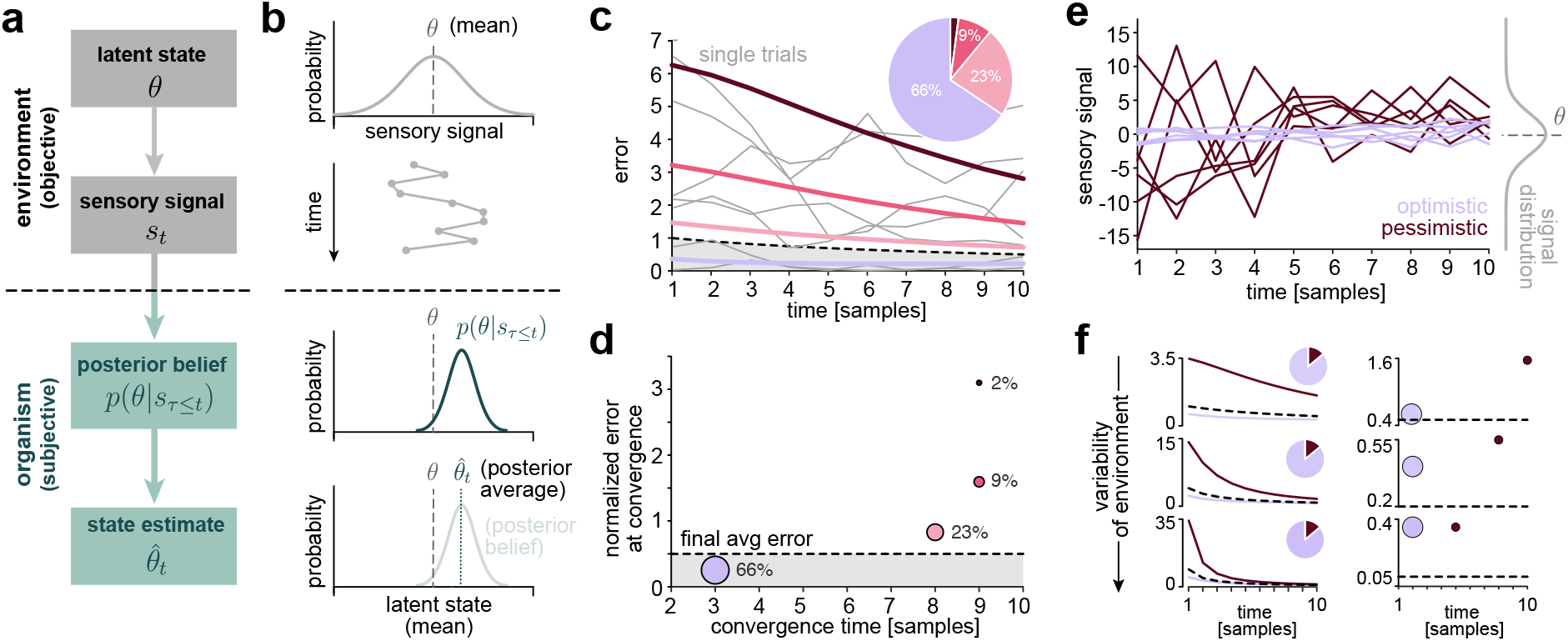
Individual inference trajectories exhibit different patterns of error. **a)** In inference tasks, an organism must estimate a latent state of the environment *θ* from available sensory stimuli *s*_*t*_. Here, we focus on scenarios where the state of the environment is unchanging in time. **b)** To perform optimal inference, an ideal observer builds a posterior belief *p*(*θ*|*s*_*τ*≤*t*_) about the underlying latent state *θ* of the environment (e.g., the mean of the distribution of sensory stimuli) from incoming samples *s*_*t*_. This belief can then be used to construct a point estimate 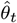 of the latent state that can be used to guide an appropriate action. **c)** The average error in the observer’s estimate (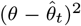black dashed line) converges slowly. However, individual trials exhibit dynamics that can deviate strongly from the mean (thin gray lines). When clustered based on their error dynamics (blue and red lines), different clusters exhibit error that is higher (red) or lower (blue) than the mean. The pie chart depicts the relative size of the error trajectory clusters. **d)** Error clusters violate the SAT. Clusters generated by optimistic sequences converge earlier and to a lower error, while clusters generated by pessimistic sequences converge later and to higher error values. **e)** Example stimulus sequences that are more (blue) or less (red) representative of the underlying stimulus distribution (“optimistic” and “pessimistic”, respectively), and that lead to lower or higher average error (colors correspond to the blue and red clusters in panels **c** and **d**). **f)** The structured variability of individual inference trials (left column) and corresponding violations in the SAT (right column) generalize across environments with more or less variability in the underlying latent state.

As a simple illustrative example, we consider the problem of inferring the mean of a Gaussian distribution (Fig. 1b), where the observer has limited time to perform the inference, and can therefore only observe a maximum of *T*_max_ = 10 stimulus samples. On average, the error in the observer’s estimate of the mean decreases in time, obeying the SAT (Fig. 1c, black dashed line). However, a simple clustering analysis reveals that this average pattern does not capture the behavior observed on individual trials (Fig. 1c, thin gray lines). For example, a majority of trials (66%) converge much faster and to a much lower error than the average (Fig. 1c, blue), while the remaining trials (34%) converge more slowly and to higher values of error. This results in a counterintuitive violation of the SAT, where faster convergence leads to lower, rather than higher, error (Fig. 1d). This variability is caused by the stochastic nature of short stimulus sequences. Purely by chance, some of these sequences are more representative of the underlying distribution, leading to a faster convergence of the observer’s estimate and a lower inference error. We call such stimulus sequences “optimistic” (Fig. 1e, blue lines). In contrast, “pessimistic” sequences are not representative of the underlying stimulus distribution and can therefore be misleading about the value of underlying latent state (Fig. 1e, red lines). These patterns of error generalize across different inference scenarios with more or less environmental variability (parametrized here by the variance in the underlying latent state; Fig. 1f).

It is worth emphasizing that this is one of the simplest scenarios that can be used to study the dynamics of inference. Despite its simplicity, individual trials exhibit structure that can be exploited to save time and bound error. Crucially, this structure is *intrinsic* to the statistics of short stimulus sequences, and would therefore be present in more complicated inference scenarios or under different model assumptions. In the remainder of the paper, we explore how this structured variability could be exploited by animals in an inference task directly related to their survival.

### An example escape task requires rapid inference under strong time limitations

Planning and coordinating an escape is a salient example of a task that animals must solve under strong time pressures. Here, we consider an important component of rapid escape: inferring the direction of an approaching predator from noisy sensory stimuli [6, 7, 20]. We study a simplified setting in which an animal must use a stream of incoming sensory stimuli *s*_*t*_ to infer a latent state *θ* that specifies the angular direction of an approaching predator (Fig. 2a, c). We assume that this latent state can take one of *N* discrete values that represent distinct approach directions. We model the animal as an ideal observer that maintains and updates a belief about the direction of approach, as summarized by the posterior distribution *p*(*θ*|s_*τ*≤*t*_). This belief can be used both to construct an estimate 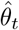 of the direction of approach, and to decide when to stop performing inference and initiate an escape (Fig. 2b,c). Importantly, while the direction of approach is drawn from one of a discrete number of possible directions, the observer’s estimate of direction 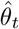 is continuous and equal to the posterior average. This estimation minimizes the squared inference error 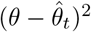 [21].

**Figure 2:**
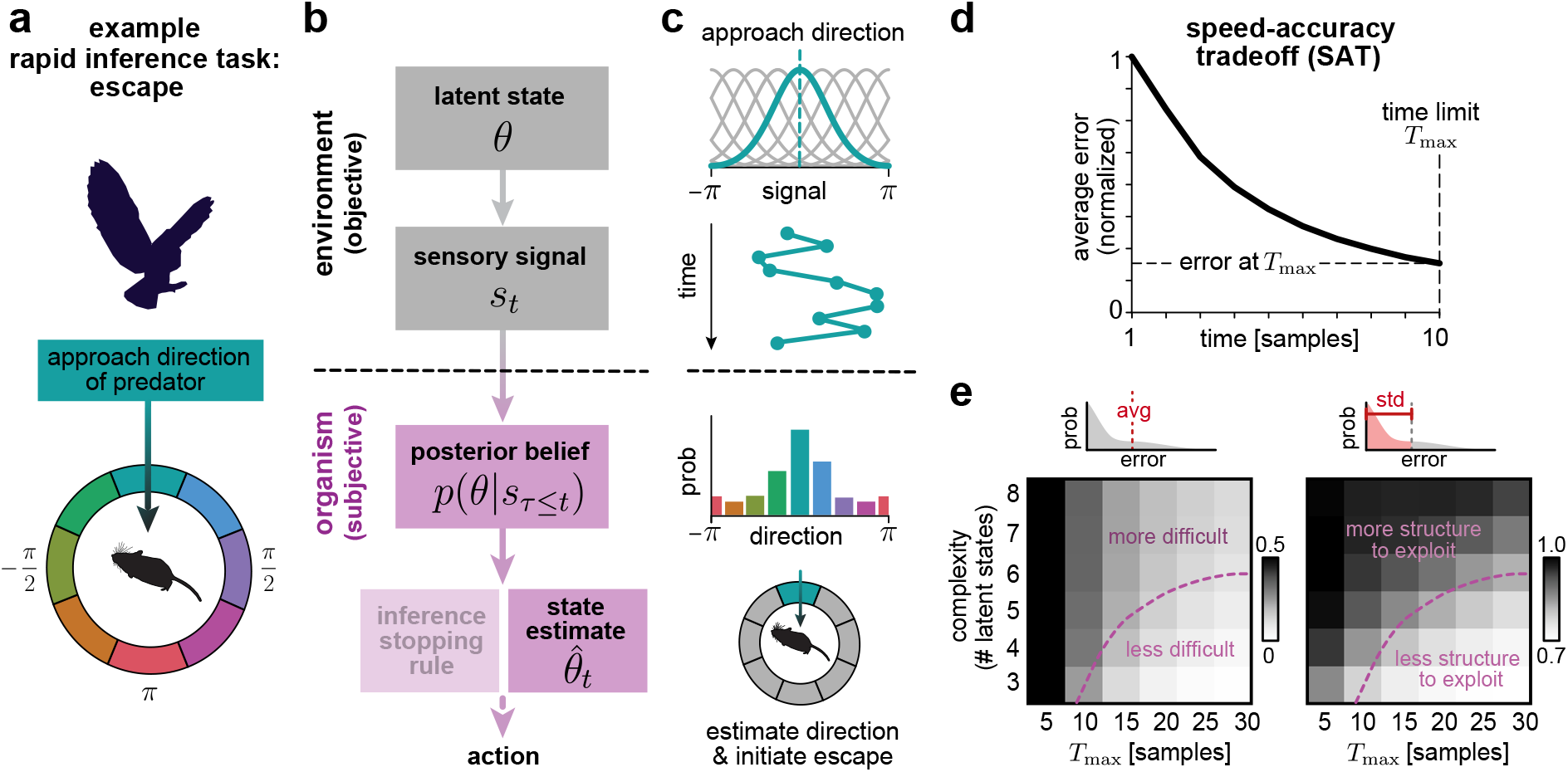
An example inference task underlies rapid escape. **a)** Example inference task: an animal infers the direction of an approaching predator to guide an effective escape. **b)** To perform inference, an ideal observer builds a posterior belief *p*(*θ*|*s*_*τ*≤*t*_) about an underlying latent state *θ* of the environment from incoming sensory stimuli *s*_*t*_. This belief can then be used to construct a point estimate 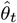 of the latent state that can be used to guide an appropriate action. Under time constraints, the observer must additionally decide *when* to stop performing inference and initiate the action. **c)** In the example shown in panel **a**, the latent state specifies the direction of approach and parametrizes the distribution of incoming sensory stimuli. These stimuli are used to infer the posterior probability distribution of different approach directions, which can in turn be used to estimate the approach direction and initiate an escape in the opposite direction. **d)** On average, longer times lead to more accurate inferences. According to this well-known speed-accuracy tradeoff (SAT), an ideal observer should use all of its available time, up to a time limit *T*_max_, to make the most accurate inference. **e)** Left: stronger time limitations (shorter *T*_max_) and more complex inferences (more latent states) lead to higher inference errors on average (“more difficult” region of heatmap). Color denotes the average error at *t* = *T*_max_, normalized by the initial error at *t* = 1. Right: under “more difficult” scenarios, there is a broader distribution of errors below the mean, and thus more structure to be exploited by an ideal observer. Color denotes the standard deviation of all errors below the mean, normalized by the standard deviation of the full error distribution.

As in the previous section, we constrain the inference period to a maximum duration *T*_max_, which limits the number of stimulus samples that can be collected by the observer. Within this limit, the trial-averaged inference error again obeys the SAT, with longer inference times leading to lower average errors (Fig. 2d). This curve suggests that the observer should continue collecting stimulus samples up until the time limit *T*_max_, since only then will the average error be minimal. Under this strategy, the final average error depends not only on the time limit *T*_max_, but also on the complexity of the inference task. In the scenario considered here, this complexity is specified by the number of latent states *N* (i.e., the number of possible directions of approach), given a fixed level of stimulus variability. More complex inference tasks require longer inference times to reach the same final error (Fig. 2e, left), and are thus more difficult under strong time pressures. While a natural solution is to wait longer to acquire more sensory stimuli and further reduce average error, this is not possible under hard time constraints. In these scenarios, although the *average* error might be too high to permit accurate decisions, the trial-specific error can be low (Fig. 2e, right), consistent with what we observed in the simpler scenario shown in Fig. 1c). This suggests that the observer could benefit from estimating its own error in order to guide appropriate actions. Moreover, if this error is likely to be low, any further reduction in inference time could be used to coordinate more precise actions. Below, we describe strategies for reducing inference time while bounding inference errors, even in seemingly challenging circumstances.

### The statistics of individual stimulus sequences differentiate patterns of inference error

Consistent with the simpler inference task considered in Fig. 1, we find that individual inference trajectories exhibit diverse patterns of error that decrease rapidly on some trials but increase on others (Fig. 3a). Distinguishing between these scenarios is crucial for performing critical tasks such as escape planning, where survival depends not on trial-averaged performance, but rather on maintaining performance above a certain threshold in individual trials. Clustering error trajectories across trials again revealed a diversity of error dynamics, with the largest cluster containing trials in which the inference error rapidly drops below the trial average (Fig. 3b). The remaining clusters exhibit errors that significantly exceed the trial average; one cluster, for example, contains trials in which the inference error is high even at the time limit (Fig. 3b, yellow line).

**Figure 3:**
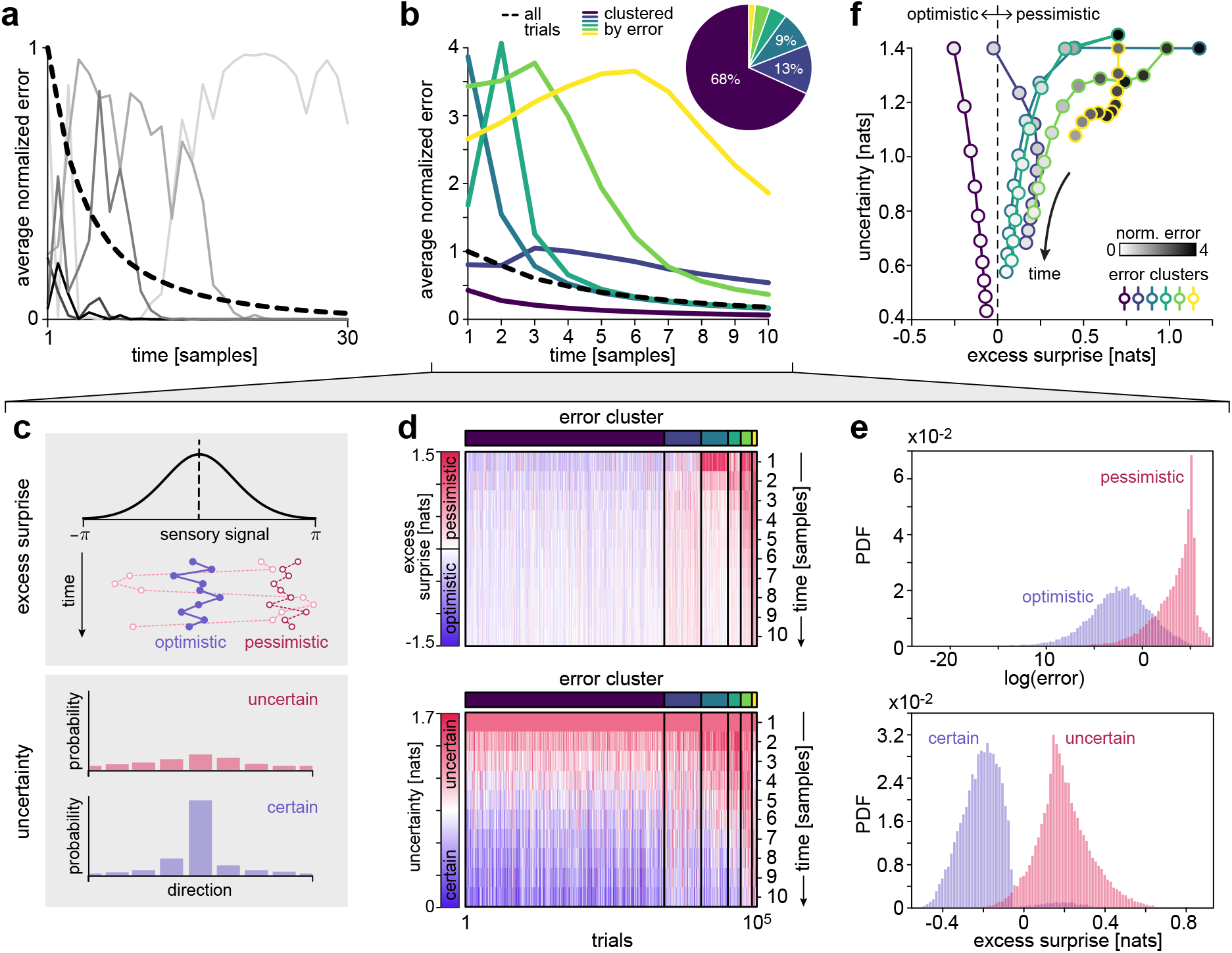
Excess surprise and uncertainty differentiate patterns of inference error. **a)** Individual trials exhibit a diversity of error dynamics. **b)** When clustered by error, different trials exhibit different patterns of average error that can violate the SAT. **c)** Schematics illustrating excess surprise (upper) and uncertainty (lower). **d)** Upper: Inference trajectories that exhibit similar patterns of error arise from stimulus sequences that exhibit similar patterns of excess surprise. These “optimistic” and “pessimistic” sequences are more or less surprising than expected on average, and lead to lower and higher average error, respectively. Lower: An ideal observer does not have access to the excess surprise of incoming stimuli, but can instead compute its own uncertainty about the underlying latent state that generated those samples. **e)** Upper: optimistic and pessimistic sequences generate distributions of lower versus higher error (blue and red distributions, respectively; shown for the top and bottom quartiles of excess surprise computed at *T*_max_ = 10). Lower: observers that were more or less certain encountered stimulus sequences with lower versus higher excess surprise (blue and red distributions, respectively; shown for the top and bottom quartiles of uncertainty computed at *T*_max_ = 10). **f)** In the limit of long inference times, uncertainty and excess surprise tend to zero, averaged within each of the clusters identified in panel **b** (line colors). At short times, both quantities can deviate from zero; these deviations correlate with high versus low error (marker fill colors). Thus, under time constraints, uncertainty can provide information about the excess surprise of incoming stimulus sequences, and, by consequence, the expected error.

As described earlier, this diversity of single trial dynamics arises from variability in stimulus sequences. Intuitively, stimulus sequences that are highly probable under the true latent state should rapidly yield an accurate inference. In contrast, stimulus sequences that are unlikely under the true latent state can be ambiguous and can lead to inaccurate inferences. To quantify this intuition, we define the *excess surprise S*_*e*_ of a stimulus sequence *s*_*τ*≤*t*_:

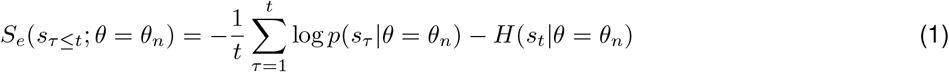

where *H*(*s*_*t*_ |*θ* = *θ*_*n*_) = −∫*p*(*s*_*t*_ *θ* = *θ*_*n*_) log *p*(*s*_*t*_ *θ* = *θ*_*n*_)*ds*_*t*_ is the expected surprise, or entropy, of the stimulus distribution given the true value *θ*_*n*_ of the latent state *θ*. Excess surprise measures the difference between the average surprise of the observed stimulus sequence and the expected surprise of the full distribution. A negative value indicates that the stimulus sequence was less surprising, and thus more probable, than expected on average (Fig. 3c, blue points); these are the “optimistic” stimulus sequences that unambiguously inform the observer about the true latent state that generated the sequence. In contrast, a positive value of excess surprise indicates that the stimulus sequence was more surprising, and thus less probable, than expected on average (Fig. 3c, red points); these are the “pessimistic” sequences that can mislead the observer to form an incorrect belief about the true latent state. We note that excess surprise bears conceptual similarity to the notion of typicality in information theory [22]. Because it measures deviations from an average property of a distribution, it is also reminiscent of concepts from large deviation theory [23].

Tracking the excess surprise over time and across trials confirms this intuition (Fig. 3d, upper panel). In the limit of long stimulus sequences, the excess surprise tends to zero (i.e., the surprise converges to the average), as expected. However, on short timescales, the excess surprise captures the observed variability in the error trajectories. Those trajectories that rapidly converge to very low error values (dark blue cluster) are generated by optimistic stimulus sequences with negative excess surprise. In contrast, when the error remains high across the duration of the trial, the underlying stimulus sequences are pessimistic, with positive excess surprise. Overall, the higher the excess surprise of the observed stimulus sequence, the larger the error at the time limit (Fig. 3e, upper panel).

These deviations from the average, which occur over short timescales, contain structure that can be exploited to inform more accurate inferences under limited time. However, the relevant quantity—excess surprise—is an *objective* measure. This means that in order to evaluate the excess surprise of a given stimulus sequence, the observer would need to know the true distribution of stimuli *p*(*s*_*t*_ |*θ*), parametrized by the true value of the latent state *θ*. Because the purpose of inference is to determine the latent state *θ*, the observer cannot directly compute the excess surprise. Instead, the observer must rely on a *subjective* quantity to which it has direct access.

One such subjective quantity is the observer’s uncertainty about the latent state, which can be measured using the entropy of the posterior distribution, *H*(*θ*| *s*_*τ*≤*t*_) (Fig. 3c, lower panel; see Methods for details). As with excess surprise, uncertainty tends to zero in the limit of long stimulus sequences. On short timescales, the dynamics of uncertainty are correlated with the dynamics of excess surprise (Fig. 3d, lower and upper panels, respectively). For example, optimistic stimulus sequences belonging to the largest cluster lead to the most rapid decrease of the observer’s uncertainty; as described above, this is because they are most probable given the true latent state, and therefore quickly lead to a correct inference. In contrast, pessimistic stimulus sequences cause the observer to maintain high uncertainty across the duration of the trial. Overall, higher uncertainty is generated by stimulus sequences with higher excess surprise (Fig. 3e, lower panel), which in turn leads to higher error (Fig. 3e, upper panel).

These relationships are also observable at the level of error clusters (Fig. 3f), where average uncertainty and average excess surprise can strongly deviate from zero in a manner that correlates with average error. For example, the highest-error cluster exhibits positive excess surprise and high uncertainty that persists throughout the trial, whereas the lowest-error cluster exhibits negative excess surprise and low uncertainty (Fig. 3f, yellow and purple lines, respectively). These relationships imply that uncertainty can be exploited by the observer to indirectly assess the excess surprise of the stimulus sequence that it encounters and, as a consequence, assess its own inference error.

### An adaptive stopping rule improves performance in difficult inference scenarios

To exploit the observed structure in single-trial inference dynamics (Fig. 3), we designed a set of adaptive stopping rules that operate on the observer’s changing uncertainty about the underlying latent state (Fig. 4a). These stopping rules are parameterized by a dynamic uncertainty threshold (Fig. 4a-1); if the observer’s uncertainty drops below this threshold on a given timestep, the observer stops performing inference and commits to an action (we will refer to these as “converged” inferences). At the time limit *T*_max_, the inference stops for one of two reasons; either the observer’s uncertainty fell below the threshold at time *T*_max_ (but not before), or the observer ran out of time. These two scenarios have different implications for the actions that the observer should take conditioned on this inference; we will first discuss the inference dynamics, and return later to the component of action selection.

**Figure 4:**
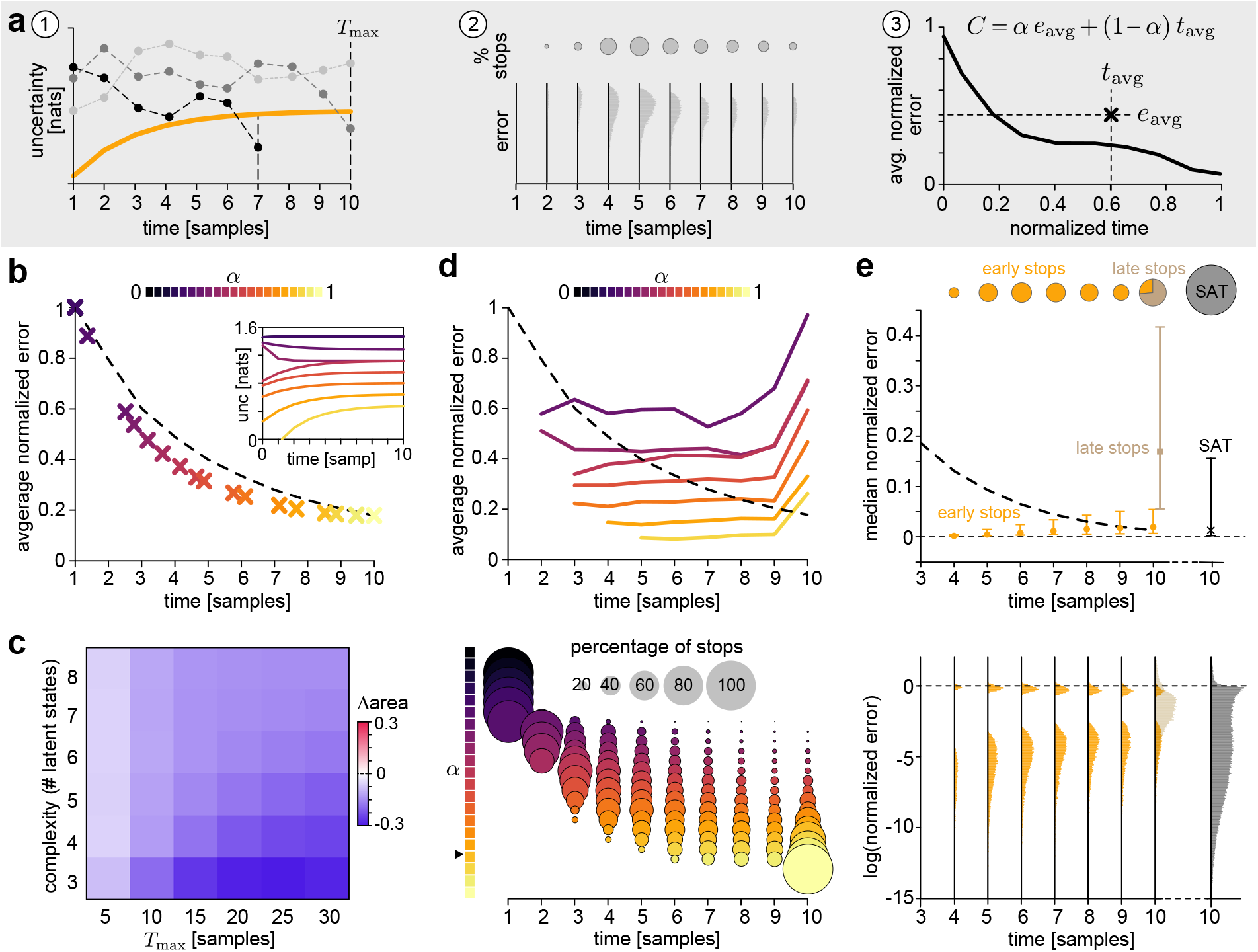
An adaptive stopping rule exploits predictable patterns of error. **a)** Schematic of adaptive stopping criteria. (1) We design an adaptive stopping rule, captured by a time-dependent uncertainty threshold, that is used to terminate the inference process up until a maximum time limit of *T*_max_. If the observer’s uncertainty drops below this threshold before *T*_max_, the inference process terminates early. At *t* = *T*_max_, the inference process can be terminated either because the observer’s uncertainty dropped below the threshold (“early” inferences), or because the observer reached the time limit (“late” inferences). (2) A given uncertainty threshold will generate a distribution of stopping times, and a distribution of errors at each stopping time. (3) We optimize uncertainty thresholds to minimize a cost *C*(*α*) that weighs the average inference error against the average inference time. Low versus high values of *α* prioritize fast versus accurate inference, respectively. **b-e)** Performance using adaptive stopping rules optimized for *N* = 5 classes and *T*_max_ = 10 samples. **b)** Higher values of *α* lead to lower average errors and higher average stopping times (lower right panel); for a given average stopping time, the average error achieved by the adaptive uncertainty threshold (different colored x’s) is lower than that achieved by the classic SAT (black dashed line). The corresponding optimal uncertainty thresholds are shown in the inset. **c)** Fractional change in area under the error-time curve, compared between the adaptive strategy and the standard SAT (Methods). Blue indicates a reduction in area, and thus an improvement in either the average error achieved at a fixed time, or the average time required to achieve a fixed error. **d)** Upper: Adaptive stopping rules generate error trajectories that violate the SAT; average error trajectories are nearly flat for *t < T*_max_, and they increase abruptly at *t* = *T*_max_. Lower: Different values of *α* generate different distributions of stopping times; higher values of *α* generate distributions that are shifted toward *T*_max_. **e)** Upper: median inference errors and inter-quartile ranges for the adaptive strategy, separated into early stops (i.e., those trajectories that converged before or at *t* = *T*_max_) versus late stops (i.e., those that did not converge). Shown for *α* = 0.8 (orange) and compared to the SAT (black). Lower: corresponding distributions of inference errors for *α* = 0.8. The adaptive stopping rule generates a bimodal distribution of errors; a small set of high errors correspond to cases in which the observer is certain but of the wrong latent state; the large bulk of low errors correspond to cases in which the observer is certain of the correct latent state. This distribution is similar for the set of early inference trajectories that terminated at different stopping times (orange distributions); in contrast, the set of late inference trajectories that did not drop below the uncertainty threshold during the time *t ≤ T*_max_ exhibit a different distribution of errors (brown). For comparison, the distribution of errors generated by the SAT at *t* = *T*_max_ is shown in black.

The adaptive stopping rules lead to a *distribution* of stopping times, each of which produces a distribution of final errors (Fig. 4a-2; upper and lower panels, respectively). This, in turn, leads to a trajectory of average error than can in principle violate the SAT (Fig. 4a-3). We use the distributions of stopping times and final errors to compute the average stopping time *t*_avg_ and average error *e*_avg_, given a particular instantiation of the uncertainty threshold. We then use these average quantities to optimize the parameters of the uncertainty threshold by minimizing the following cost function:

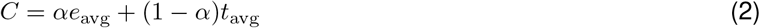

where *α* ∈ [0, 1] is a parameter that controls whether the observer prioritizes the speed (*α* near 0) or accuracy (*α* near 1) of inference.

When optimized in this way, the adaptive stopping rules generate lower average error than the SAT for the same average stopping time (Fig. 4b); this improvement is consistent across different time constraints and inference tasks (Fig. 4c). However, the underlying trajectories of error and distributions of stopping times are qualitatively different than the SAT (Fig. 4d); for *α* near 0, the observer stops quickly to prioritize inference speed; for *α* near 1, the observer tends to wait longer to prioritize inference accuracy (Fig. 4d; lower panel). When averaged at each stopping time, the resulting error trajectories strongly violate the SAT, maintaining near-constant errors until the time limit, when the error increases (Fig. 4d, upper panel).

This behavior is even more apparent when we decompose inference trajectories into those that fell below the uncertainty threshold before or at the time limit (“early stops”) versus those that were forced to stop at the time limit (“late stops”) (Fig. 4e, shown for *α* = 0.8). Early stops generate low errors whose distribution is consistent across time; for this particular value of *α*, the distribution is bimodal, corresponding to scenarios in which the observer was sufficiently certain to stop the inference process but was either correct (low error) or incorrect (high error) in its estimation of the latent state. Late stops, in which the uncertainty did not drop below the threshold and the observer was forced by the time limit to stop the inference process, show a qualitatively different distribution of error. Thus, these different types of stops implicitly carry information about the underlying error, and could thus be used to guide distinct types of actions.

### Adaptive stopping rules exploit the statistics of individual inferences

To gain a more mechanistic understanding of this adaptive stopping rule, we examined how it altered the joint distribution of error and excess surprise over the course of inference (Fig. 5a). The full distribution, computed across all trials, spans a wide range of errors and values of excess surprise. The adaptive stopping rule can be viewed as subselecting a fraction of trials from this full distribution; the remaining trials then constitute the distribution that can be subselected on the next timestep (Fig. 5b). Consistent with the trial-to-trial structure observed in Fig. 3, the adaptive stopping rule—which is based only on the observer’s uncertainty—selects those times and trials with negative excess surprise. These predominantly correspond to trials with low errors, but include a small fraction of trials in which the observer was certain but of the wrong latent state, and thus had high error. As time passes and the excess surprise tends toward zero, the adaptive stopping rule selects trials with higher (but still negative) excess surprise that nevertheless generate similar distributions of error.

**Figure 5:**
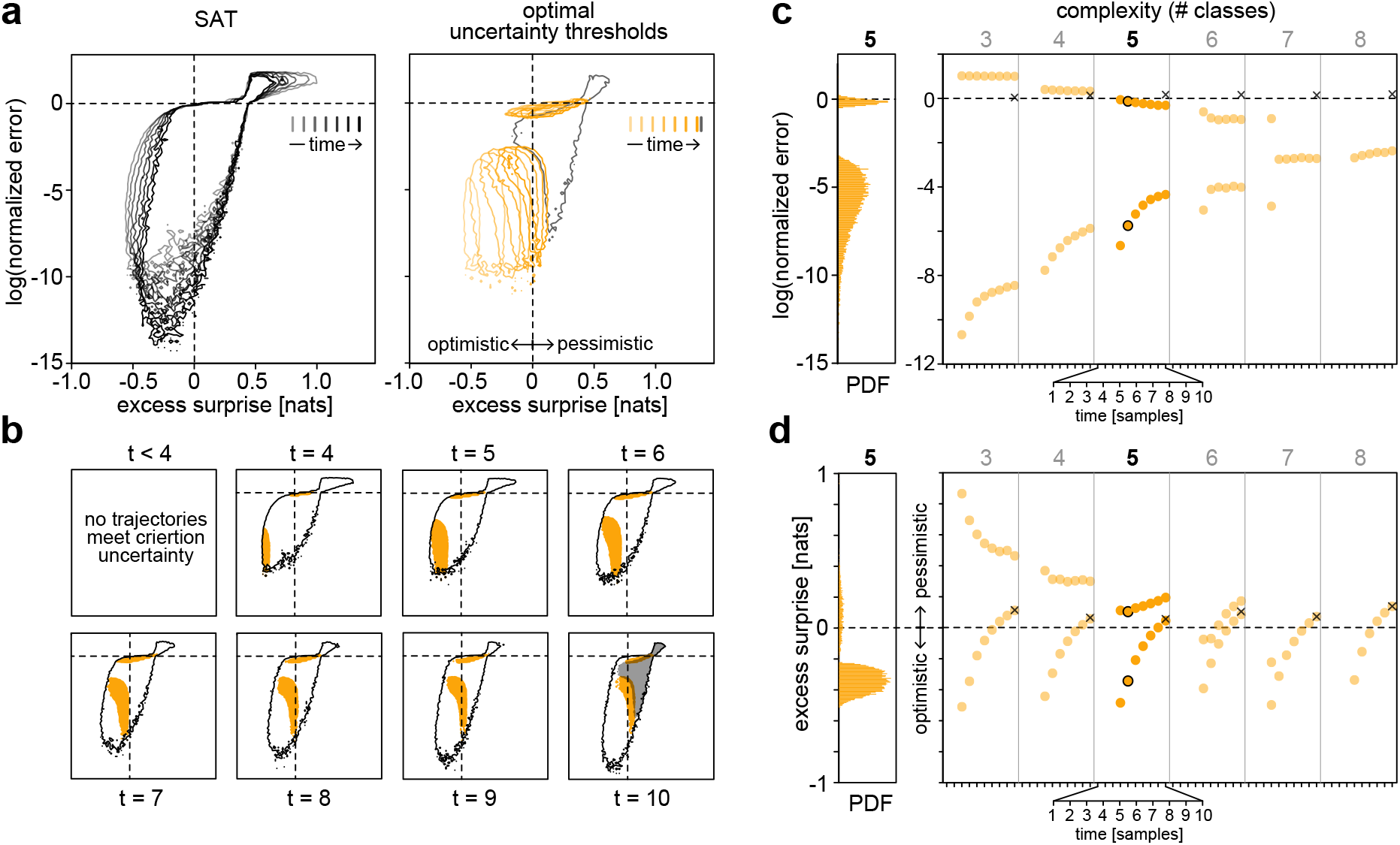
The adaptive stopping rule exploits optimistic stimulus sequences that generate low uncertainty. **a-b)** Joint distributions of error and excess surprise for different stopping rules. **a)** Left: a fixed stopping rule based on the SAT generates distributions that encompass all inference trajectories. Right: the adaptive stopping rule subselects a set of inference trajectories that fall below the uncertainty threshold at time *t*; the remaining inference trajectories contribute to distributions at times *t*^*′*^ *> t*. Early stopping times (light colors) are driven by the most optimistic stimuli that rapidly yield low uncertainty. As time goes on, the remaining inference trajectories are driven by less-optimistic stimuli. Nevertheless, the adaptive stopping rule selects from among these trajectories to achieve a similar distribution of errors. All trajectories that did not converge in a time *t* ≤*T*_max_ are forced to stop at *T*_max_; these make up the gray distribution. **b)** Same as panel **a**, but split out by different stopping times and compared between the fixed SAT rule (black outlines) and the adaptive stopping rule (filled regions). As in **a**, the distribution at *t* = *T*_max_ is made up of two different types of stops; those that that fell below the uncertainty threshold (orange), and those that did not (gray). **c)** Under the adaptive stopping rule, the distribution of errors is bimodal across inference tasks of varying complexity. This bimodality eventually collapses for sufficiently high numbers of inference classes, because there is no longer a strong separation in error between correct versus incorrect inferences at a given uncertainty level. Pairs of orange markers denote the average errors of high-versus-low error modes of the distribution, computed for early inferences. Black x’s denote the average error of late inferences. Black open circles for *N* = 5 correspond to the error distribution to the left of the panel. **d)** Average excess surprise for inference trajectories that correspond to high-versus low-error modes of the distributions shown in panel (c). Low error modes correspond to more optimistic stimulus sequences. Black circles for *N* = 5 correspond to the excess surprise distribution to the left of the panel.

These results are qualitatively consistent across inference scenarios with varying complexity (Fig. 5c,d). In simpler settings, the error distribution is strongly bimodal (Fig. 5c), corresponding to scenarios in which the observer was certain about the correct versus incorrect state, as signaled by optimistic or pessimistic stimulus sequences, respectively (Fig. 5d). As the complexity increases, it becomes more difficult to differentiate latent states, and the two modes of the distribution begin to merge. In all cases, those trials that did not converge before or at the time limit had high average error that was generated by pessimistic stimulus sequences (black x’s in Fig. 5c-d).

Together, these results confirm that the adaptive stopping rule leverages trial-to-trial variability by differentiating optimistic versus pessimistic sequences, enabling the observer to make accurate inferences in far less time.

### Adaptive stopping confers several key advantages

Adaptive observers indirectly exploit the statistical structure of very brief stimulus sequences. This strategy has multiple advantages that increase the probability of accurate inferences and, as we will show, the actions that depend on them.

First, adaptive inference rules provide subjective information about the objective level of error. If an inference does not converge within the time limit, it is highly likely that the underlying stimulus sequence is pessimistic and the inference error is large. Alternatively, if an inference does converge, it is highly likely that the underlying stimulus sequence is optimistic and the inference error is small, regardless of *when* the inference converges (Fig. 4e-f; Fig. 5a-b). This meta-knowledge is related to the notion of confidence [24] and can be exploited by the observer to plan appropriate actions. For example, in planning an escape, animals can freeze if time has run out and they remain uncertain of the direction of an approaching predator; alternatively, if their inference rapidly converges, they can plan coordinated actions under the assumption of a reliably accurate estimate of the approach direction.

Second, those inferences that converge within the time limit are substantially faster, and have substantially lower error, when compared to the “static” SAT strategy (Fig. 4g-h). This is because they exploit random fluctuations in short stimulus sequences, information that is not used if the observer merely waits a predetermined amount of time. Third, adaptive stopping improves the efficiency of the underlying sensory processing that supports inference.

Because adaptive observers terminate inference as soon as their uncertainty has dropped below threshold, they use far fewer stimuli on average to achieve the same accuracy of inference as an observer that waits for a predetermined amount of time. As a result, the underlying sensory system has fewer stimuli to encode, process, and transmit.

To concretely illustrate the advantages of adaptive stopping, we couple our ideal observer to a decision maker that can select and initiate actions, and we use this to model a scenario that mimics escape behavior in the fruit fly, *Drosophila melanogaster* (Fig. 6a). We model the fly as an agent that has limited time to infer the direction of a looming predator and execute an escape in the opposite direction. We assume that this takes places in two stages: first, the model fly has a maximum time limit of *T*_max_ to infer the approach direction from noisy stimuli that signal one of *N* latent states; here, *T*_max_ serves as a proxy for the looming speed of the predator, whereby faster looming speeds impose stronger time limitations on inference. Second, after inferring the direction of approach, the model fly must execute an escape in the opposite direction before colliding with the predator. Here, we assume that there are tradeoffs between the timing, precision, and accuracy of the escape: we assume that more precise actions require more time to execute, and that more accurate actions allow more time before a collision with the predator (heatmap in Fig. 6a; Methods). Thus, the model fly can successfully avoid a collision by slowly coordinating a precise and accurate escape away from the predator, or by quickly making an imprecise and inaccurate escape.

**Figure 6:**
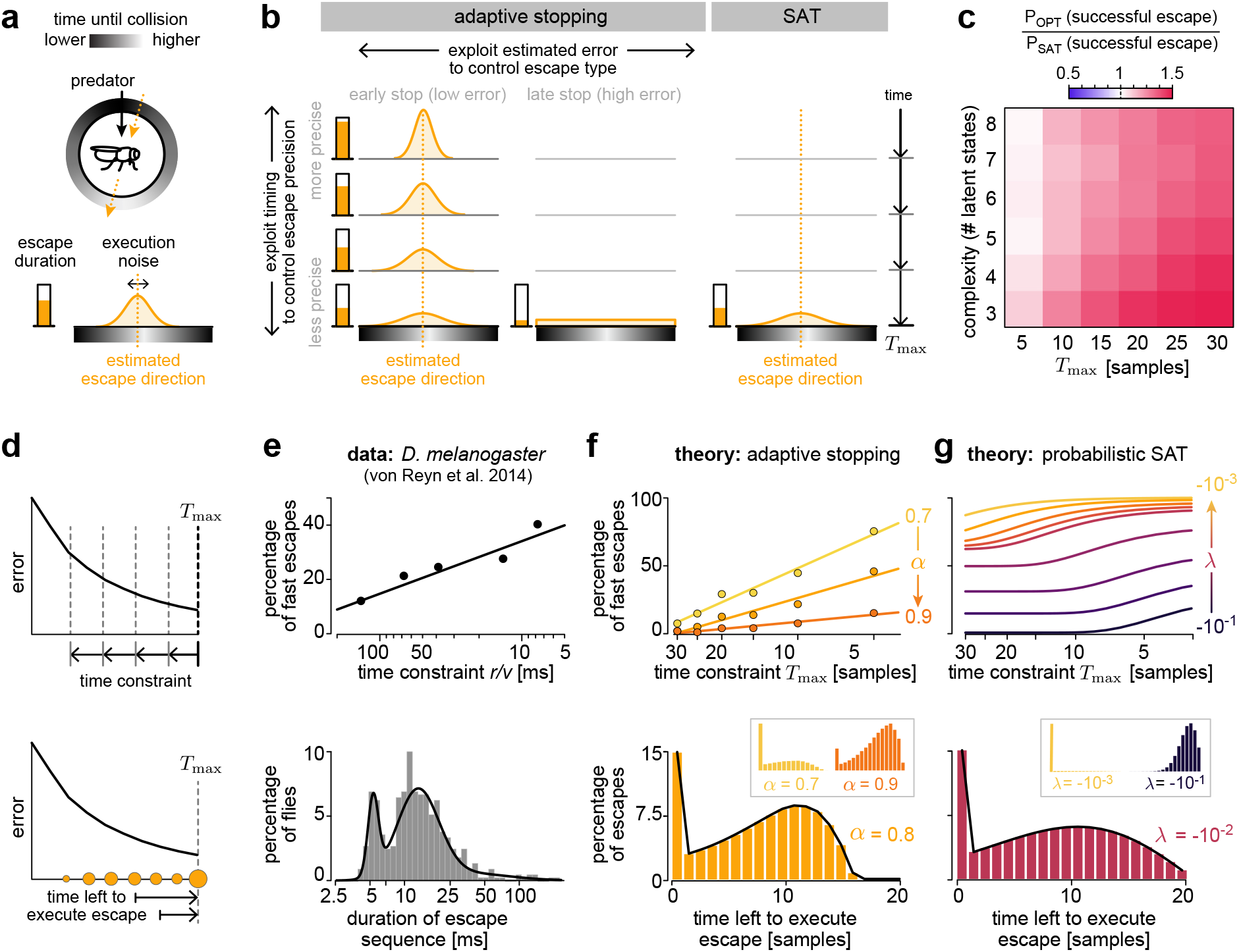
Adaptive stopping improves successful escape and reproduces qualitative features of fruit fly behavior. **a)** We consider a scenario in which a model fly (i.e., our ideal observer) must infer the direction of an approaching predator and use the inference to guide an escape in the opposite direction. The inference must be completed with *T*_max_ timesteps; the execution of the escape requires additional time and precision. We compute the probability that the escape is executed before the model fly collides with the predator; we assume that collisions happen later if the model fly escapes away from the predator (grayscale colormap), and that more precise actions—which are more likely to avoid the predator—require more time to execute (orange shaded distribution). **b)** The adaptive stopping rule (left and middle columns) can be exploited to improve the precision and type of escape (rows and columns, respectively); more rapid inferences can be used to coordinate more precise escapes, as indicated by the narrower distributions at top. Similarly, early versus late inferences can be used to trigger coordinated escapes that are centered on the estimated escape direction but are slower to execute (left column), versus last-ditch escapes that are executed in a random direction but are fast (middle column). The SAT (right column) does not permit such flexibility. **c)** Adaptive stopping improves the probability of a successful escape across a wide range of time constraints and inference tasks (red versus blue respectively denote an increase versus decrease in the probability of successful escape). **d-g)** Statistics of escape behavior compared between the fruit fly, the adaptive stopping model, and the probabilistic SAT model, measured as a function of time constraint (upper row) and escape duration (bottom row). **d)** Schematics illustrating the different factors that are compared in panels **e-g. e)** Escape behavior of *Drosophila melanogaster* in response to rapidly looming visual stimuli (reproduced from [26]). Upper: fraction of fast and imprecise escapes as a function of the visual speed of the looming stimulus. Faster looming corresponds to lower *r/v* values, where *r* is the stimulus size and *v* is the looming speed. Lower: distribution of escape durations. Modes of the distribution correspond to two escape modes—fast and imprecise escapes of short duration, and slow, deliberate escapes of long duration. **f-g)** Escape behavior predicted by the adaptive stopping model (**f**) and the probabilistic SAT model (**g**). Upper: fraction of fast escapes as a function of the time constraint *T*_max_, which limits the maximum duration of inference and serves as a proxy for the looming speed. Colors correspond to different values of the trade-off parameter *α* (**f**) or the difficulty parameter *λ* (**g**). Lower: probability distribution of escape durations, measured as the time between the initiation of the escape and the time limit *T*_max_=20 samples. Insets depict probability distributions for different parameter values.

In such a scenario, an adaptive model fly can exploit the timing and expected error in its inference to improve the probability of a successful escape (left and middles column in Fig. 6b). If its certainty drops below threshold at or before the time limit, we assume that the model fly can initiate a slow escape (left column in Fig. 6b). In this situation, the model fly believes that it has a correct estimate of the approach direction, and it can use its remaining time to plan and execute a precise escape; the earlier the inference, the more time it has to execute a sequence of actions, and the more precise the escape will be (Fig. 6b, left column; note that execution time is higher, and the execution noise is lower, when the animal is further from the time limit). Alternatively, if the model fly reaches the time limit before its estimate has converged, it can initiate a fast escape. In this situation, the model fly is uncertain about the direction of approach, its estimation error is likely high, and the time for inference has run out. Rather than using its erroneous estimate, we assume that it instead initiates a “last ditch” fast escape that is taken in a random direction but at a more rapid speed (Fig. 6b, middle column; note that execution time is shorter and the execution noise is flat, compared to a slow escape taken at the same time).

In contrast, a decision based purely on the SAT does not have information to guide different patterns of escape based on timing or error (Fig. 6b, right column). This leads to lower probability of a successful escape compared to the adaptive strategy, regardless of the time constraint or the complexity of the inference task (Fig. 6c).

### Escape behavior in *Drosophila melanogaster* exhibits signatures of adaptive stopping

The scenario described above (Fig. 6a-c) highlights some of the key advantages of using an adaptive stopping rule to inform decisions and actions (Fig. 3-5). To explore whether these advantages might be borne out in real escape behavior, we compared the properties of the underlying inference to measurements of escape behaviors in the fly. Importantly, these properties were intrinsic only to the inference process, and were independent of the model of escape decisions described above. In the fly, escape behavior has typically been studied by exposing flies to an expanding visual disc that mimics an approaching predator [8, 20, 25, 26]. Flies exhibit two different modes of escape in response to such stimuli [20]: either a slow, precise escape that is preceded by an elaborate sequence of preparatory movements, or a fast, imprecise escape without extended preparation. These two different modes are mediated by different neural pathways [26], and they require different amounts of time to execute (Fig. 6e, bottom; reproduced from [26]). The probability that the fly initiates a fast escape increases with the speed of the looming stimulus; in other words, the faster the loom, the less time there is to make a decision and execute a series of preparatory actions, and the more likely the fly will initiate a fast and imprecise escape (Fig. 6e, top; reproduced from [26]). Distinct modes of escape behavior have also been observed in other species, including in fish, where they are controlled by the Mauthner cell circuit [27].

As illustrated above, our theory naturally produces two distinct types of inference that could underlie these different modes of escape: “certain” inferences that fall below the uncertainty threshold within the time limit and can be used to guide slow, precise escapes, and “uncertain” inferences that could be used to trigger a fast, imprecise escape. We find that the probability of initiating a fast escape scales linearly with the logarithm of the time limit in a manner that resembles fly behavior (compare the upper panels of Fig. 6f and 6e). Furthermore, we observe a bimodal distribution of escape durations, which in our model corresponds to the remaining time between the decision to escape and the time limit *T*_max_ (compare the lower panels of Fig. 6b and 6a). Both of these features are consistent across a range of *α* values. We note that the time remaining for inference and the time remaining for action are partially confounded in existing experimental data; future experiments could disentangle their individual impact on escape behavior.

In contrast, these two distinct modes of escape behavior cannot easily be explained as a manifestation of the SAT. The SAT suggests that later actions—which would occur after a longer collection of sensory stimuli—should in principle be more accurate than earlier actions. Thus, according to the SAT, the only rational strategy would be to wait to initiate an escape until the last possible moment, when the fly’s inferences are most likely to be accurate. This would give rise to highly stereotyped behavior, in contrast to what is observed in real flies. To permit a more rigorous comparison between fly behavior and the SAT, we derived a family of probabilistic stopping rules that directly exploit the SAT (Methods). At each timepoint up until *T*_max_, an the model fly probabilistically decides whether to continue performing inference, or whether to stop the inference process and initiate an escape. We assume the decision to continue performing inference is informed solely by the SAT curve; the higher the average error at a given timepoint, the more likely the model fly is to continue performing inference. To capture a variety of inference scenarios, we model this SAT curve as an exponential function parameterized by *λ ≤* 0. When *λ* is close to 0, the inference task is difficult, and the animal favors performing inference; this tends to force a large fraction of fast escapes at the time limit. For *λ* large and negative, the inference task is easy, and the animal favors initiating a slow and early escape. In contrast to the adaptive stopping rule, we find that this probabilistic SAT rule does not reproduce the behavior of real flies (Fig. 6b-c); the fraction of fast escapes does not scale linearly with the logarithm of the time limit for any value of *λ* (Fig. 6c, upper), and the resulting distribution of escape durations is bimodal for only a very narrow range of *λ* values (Fig. 6c, lower).

Together, these results suggest that escape decisions in the fly might exploit principles of rapid inference similiar to the adaptive strategy described here. This strategy would naturally give rise to the types of variability observed in this highly optimized behavior.

## DISCUSSION

To survive, animals are faced with the daunting task of making life-threatening inferences based on a limited set of noisy stimuli. In this work, we identified statistical regularities that could be exploited in such dire situations. Our key observation is that on very short timescales, inference errors can strongly deviate from their average behavior. We demonstrated that the structure of such deviations could be used by an ideal observer to decide when to stop collecting stimuli in order to increase the speed of inference and bound the resulting error. We further showed that the behavior of such observers can locally deviate from the classical speed-accuracy tradeoff (SAT)—a pattern that emerges on average. Finally, we showed how this adaptive strategy could be used to guide more effective escape behavior, and identified similarities with escape behavior in the fruit fly *D. melanogaster*.

This work highlights the relevance of maintaining not only point estimates of behaviorally relevant quantities, but also the “meta knowledge” captured by the observer’s perceptual uncertainty about the accuracy of those point estimates. The dynamics of the observer’s belief can provide useful information beyond the observer’s “best guess” about the state of the environment. Our observer uses this meta knowledge to decide when to stop gathering sensory stimuli and commit to an action. The problem of deciding when to stop collecting data during dynamic inference is often studied in the context of Bayesian optimal stopping [28] or sequential decision making [29]. These problems are typically solved through backwards induction and dynamic programming [30], and often require specifying a cost to taking an action [31]. Here, we consider an alternative approach that exploits the idiosyncratic properties of very brief sequences of observations, without invoking the downstream cost of taking actions based on those observations. Deviations of random variables from their asymptotic behavior fall within the purview of large deviation theory [23], but are typically analyzed using objective measures that are not available to an animal. Here, we contend with an animal’s need to make subjective inferences of such objective quantities. Moving forward, a synthesis of dynamic Bayesian inference and large deviation theory could be used to devise more sophisticated approaches for performing rapid inferences with very limited samples.

Our inference model captured only very basic aspects of rapid inference and escape planning in animals. Here, we assumed that the observer knows how much time is available for performing inference, and it can optimize its inference subject to this time limit. In real-world settings, the observer must infer this time limit from the same incoming stimuli that it uses to estimate other environmental properties, such as the direction of an approaching predator. Moreover, real-world inference must be performed using complex stimuli, such as images [32], sounds [33], or smells [16], that evolve over time and according to multiple interacting latent states. For example, an animal might need to infer the shortest path towards its shelter, together with the evolving trajectory of the predator [34].

However, despite the simplicity of our scenario, we identified short-timescale structure that could be exploited to increase inference speed; we therefore expect complex scenarios to exhibit even richer structure that could be exploited to further optimize rapid inference and planning. This simple scenario is the first normative perspective to capture several qualitative features of escape behavior in the fruit fly, including (i) the existence of two different modes of escape based on two qualitatively different outcomes of inference, (ii) the relative propensity of each mode based on the time limitations of inference, and (iii) the distribution of escape durations made within each mode. If flies indeed exploit the short-timescale structure of incoming stimuli, we would additionally expect that the error of their inferences would be low for all long-mode escapes, regardless of whether these escapes were initiated earlier versus later in time. This would suggest that observed variability in the accuracy of the escape arises from the extent of preparatory movement, and not from the accuracy of the inference that preceded the movement. This highlights the challenge inherent in studying the dynamics of rapid inference and escape planning, namely that the time for performing inferences and for executing actions are both strongly limited. Under our interpretation, flies perform fast and inaccurate escapes because their perceptual estimate never converged, and thus they were left without any remaining time to execute a long, deliberate escape. One possibility is that the time allotted by the animal to plan the escape is shorter than the time to infer and act together. New experiments will be necessary in order to dissociate these and other factors. For example, it has been demonstrated that when planning escape routes, mice use heuristics that modularize the space of parameters that describe the escape path [35]. The theory presented here could be broadened to make predictions about specific features of actions that could result from rapid inferences.

Our approach is complementary to a long tradition in computational neuroscience of using using drift-diffusion models (DDMs) to study the dynamics of decision making. Within this class of models, decision making can be understood as a specific form of categorical inference in which an observer must discern one of several (typically two) alternative distributions of the data [36]. These models reveal rich and diverse phenomenology (see e.g. [10]) and can easily be augmented with features such as dynamic boundaries or urgency signals that allow the observer to control the expected duration of inference [31, 37]. Moreover, the parameters of DDMs can be mapped onto neural quantities that can be directly observed in experiments [38]. In this work, we take a more general theoretical approach that can be more naturally extended to a broader range of scenarios, but comes at the cost of immediate points of comparison to neural data. The principles described here apply to any Bayesian inference scenario and are therefore not limited to the categorical inferences solved by DDMs. As such, they can easily be extended to any problem in which an observer must make a rapid decision based on an evolving posterior belief about a behaviorally-relevant state of the environment, including multiple continuous latent states and stimuli. Moreover, we do not need to augment the ideal observer in order to control the duration of inference; rather, we ask how the observer could use its own posterior belief to exploit structure *intrinsic* to individual stimulus sequences, and we analyze the patterns of timing and error that would result from such a strategy.

The cost of this generality is that this work cannot, in its current form, be readily and directly mapped onto specific neural quantities, such as the firing rates of neurons during decision making under uncertainty [10, 14, 37]. Several lines of existing work have proposed neural implementations of decision making that follow the SAT [9, 10]. Yet other approaches consider neural representations of uncertainty that could underlie such probabilistic computations [39]. In flies, the neural basis of escape behavior has been mapped onto different descending pathways that mediate short-versus long-mode escapes [26]. Neural pathways dedicated to the control of escape behaviors are also known in mice [32] and fish [27, 40]. Whether and how these pathways might be shaped by ongoing inferences in a manner that explains individual variability in behavior remains an interesting direction for future research.

### Outlook

To survive, organisms must often exploit every available source of information. In this work, we demonstrate that brief stimulus sequences, although seemingly random, contain enough information to allow organisms to tip the scales in favor of increasing speed during critical inferences that may decide life or death in a fraction of a second.

## ACKNOWLEDGEMENTS

We thank Gwyneth Card for useful discussions about escape behavior in flies. AMH was funded by the Howard Hughes Medical Institute.

## METHODS

### Time-constrained inference of the Gaussian mean

To illustrate structured variability on individual inference trials, we consider a simple Bayesian model of Gaussian mean inference, fully described in [41]. In brief, the observer uses stimuli *s*_*t*_ that follow a Gaussian distribution with an unobserved mean *θ* = 0 and known variance *σ*^2^ = 9. The posterior over the latent mean *p*(*θ*|*s*_*τ*≤*t*_) is also a Gaussian distribution, whose mean 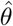 is the minimal squared-error estimate of *θ*. We simulated K = 1, 000, 000 inference trials, and we clustered the error trajectories using the standard k-means algorithm. We determined the convergence time of a trajectory to be the time point where the error is within 20% of the final error of that trajectory.

To consider environments with different variability, we considered three different values of the variance 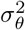 of the latent mean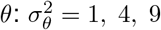.

### Time-constrained inference of the direction of an approaching predator

We consider a scenario in which an ideal observer must infer the direction of an approaching predator from one of *N* possible angles of approach *θ* ∈ {*θ*_1_, …, *θ*_*N*_} (Fig 1a-b). To infer this direction, the observer relies on a brief sequence of noisy sensory stimuli 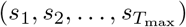, where *T*_max_ limits the total inference time. Sensory stimuli are drawn from a von Mises distribution; i.e., *p*(*s*_*t*_|*θ*) = vonMises(*θ, κ*), where *κ* = 1 is the noise level (Fig 1c). At each time step *t*, the observer updates the posterior distribution *p*(*θ*|*s*_1_, …, *s*_*t*_) and computes the estimated direction 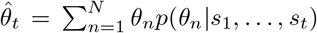. We compute the inference error at time *t* as 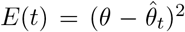 to compare across different inference scenarios, we compute the normalized error *Ē* (*t*) = *E*(*t*)/ ⟨*E*(1)⟩ _*K*_, where the normalization is performed relative to the average initial error at time *t* = 1, computed over a set of *K* inference trajectories.

#### Quantification of error dynamics

We performed inference using *N* = [3, 4, 5, 6, 7, 8] latent states. For each number of latent states, we simulated *K* = 100, 000 trials of inference over *T* = 30 timesteps. Unless otherwise stated, all results in the main text were shown for *N* = 5 latent states using a maximum inference time of *T*_max_ = 10. In Figure 3, we used k-means clustering to cluster error trajectories into 6 clusters; we chose this number for illustration, but our results were qualitatively consistent for different numbers of clusters. We used Eq. 1 to compute the excess surprise of a given stimulus sequence up to and including time *t*. To compute uncertainty, we used the entropy of the posterior at time *t*:

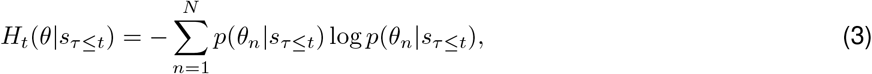

where *s*_*τ*≤*t*_ = *s*_1_, …, *s*_*t*_ is the specific sequence of sensory stimuli up to and including time *t*. Note that by notation *H*_*t*_(*θ*| *s*_*τ*≤*t*_) we mean entropy of the posterior at a time-point *t*, not conditional entropy of *θ* given *t* samples long sequence of stimuli. When analyzing distributions of uncertainty and excess surprise, as shown in Fig. 3e, we split the inference trajectories into the bottom and top quartiles of uncertainty (Fig. 3e, lower panel) or excess surprise (Fig. 3e, upper panel), computed at time *t* = *T*_max_ = 10.

#### Optimization of adaptive stopping rules

We parameterized uncertainty thresholds using the following functional form: 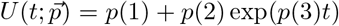. We optimized the parameters 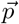 to minimize the cost function given in Eq. 2, using 21 different values of *α* evenly spaced between and including 0 and 1. Given a fixed ensemble of *K* inference trajectories, each parameter setting will cause a fraction 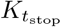 of these trajectories to fall below threshold at a time *t*_stop_ ≤ *T*_max_. Any trajectories that did not fall below threshold before or at time *T*_max_ were stopped at *T*_max_. Fig. 4b shows the average normalized error *e*_avg_ and stopping time *t*_avg_ for different *α*; these are the values that are optimized in Eq. 2 for a given *α*. To measure the improvement in this strategy compared to the standard SAT (Fig. 4c), we interpolated the set of values {(*e*_avg_(*α*), *t*_avg_(*α*))} at timepoints *t*_interp_ = [1, …, *T*_max_] to compute a version of an adaptive speed-accuracy tradeoff; we then measured the difference in the area under this curve, relative to the non-adaptive SAT: Δarea = ⟨SAT_adaptive_ −SAT_non−adaptive_⟩ / ⟨SAT_non−adaptive_⟩.

To examine violations in the speed-accuracy tradeoff, we used the distribution of trajectories that stopped at each value of *t*_stop_ to compute the average normalized error as a function of time, 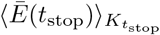. These curves are shown in Fig. 4d, together with the corresponding distributions of stopping times. We then decomposed inference trajectories into those trajectories that converged before or at *t* = *T*_max_ (“early stops”), and those that did not (“late stops”). Within each group, we computed the median, 25%, and 75% quantiles of the distribution of normalized error at each stopping time (upper panel of Fig. 4e); the full error distributions are shown in the lower panel of Fig. 4e.

#### Simplified model of escape via adaptive stopping

We built a simple model to illustrate the computational advantages of adaptive stopping. We note that due to multiple factors that determine successful escape (the duration of the escape, the timing of inference, and the accuracy of the latent state estimate), this scenario would be difficult to model using standard methods of Bayesian decision theory. Analogously to the results presented in Fig. 4, we simulated an ideal observer that estimates the direction of an approaching predator, and uses this estimate to initiate an escape in the opposite direction. The ideal observer uses an adaptive uncertainty threshold to determine when to stop performing inference and initiate an escape; if the inference converges before or at the time limit *t* = *T*_max_ (“early stops”), the observer initiates a slow escape; otherwise (“late stops”), the observer initiates a fast escape. We assume that slow escapes are more precise but require longer time to execute; in contrast, fast escapes are imprecise but are fast to execute. We model this by assuming that fast escapes are executed in a random direction and cost a time Δ*t*_fast_ ∼*P*(Δ*t*; *λ*_fast_), where we take *P*(*x*; *λ*) = exp(™*x/λ*)*/λ*, and *λ*_fast_ = *C*_fast_ = 0.02. We assume that slow escapes cost a time

Δ*t*_slow_ ∼ exp(Δ*t*; *λ*_slow_(*t*_stop_)). We assume that *λ*_slow_ = *T*_max_ ™ *t*_stop_ + *C*_slow_; i.e., the average escape times scales with the time remaining between the time at which the inference converged, and the maximum time limit. We take *C*_slow_ = 0.1, such that slow escapes at *t* = *T*_max_ cost more time than fast escapes.

In addition to differences in duration, we assume that slow and fast escapes differ in their precision. In contrast to fast escapes, which are executed in a random direction, we assume that slow escapes are initiated at the direction specified by the inference process, but that they are corrupted by execution noise whose variance scales with *T*_max_ ™ *t*_stop_. In other words, for early inferences that stop well before the time limit, the observer has a long amount of time to coordinate an escape, and the execution noise is low; for late inferences that stop near the time limit, the observer has little time to coordinate an escape, and the execution noise is high. We use this to assume that the executed escape direction is *θ*_exec_ ∼ vonMises 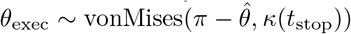, where 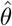 is the observer’s estimate of the direction of predator approach. We take *κ*(*t*_stop_) = 1 + *T*_max_ ™ *t*_stop_; i.e., the variance increases as *t*_stop_ approaches *T*_max_.

Given an executed escape direction *θ*_exec_ and duration *t*_exec_, we compute the probability that the escape is executed within a time limit that depends on orientation from the predator: *P*(successful escape) ≡ *P*(*t*_exec_ *< t*_thresh_(*θ*_exec_)), where *t*_thresh_(*θ*_exec_) = vonMises_*s*_(*π ™ θ ™ θ*_exec_, *κ*_thresh_); in other words, *t*_thresh_ is largest (and thus there is the highest probability of escape) when the escape is executed in the opposite direction as the predator (i.e., when *θ*_exec_ = *π* −*θ*). Here, vonMises_*s*_ denotes that the von Mises function is scaled by its maximum value, such that *t*_thresh_(*π™ θ*) = 1. In practice, we use the joint distribution of escape directions and escape times to compute *P*(successful escape) for an individual inference trajectory; we then average this probability across *K* = 100, 000 inference trajectories for a given time limit and inference task. Fig. 6c compares the performance of this model to an analogous SAT version that only uses slow escapes at time *t* = *T*_max_.

#### Comparisons to data

We compared models of dynamic inference to the behavior of *Drosophila melanogaster* published in [26]. We used two behavioral features as a basis for comparison: i) the probability of eliciting a fast escape as a function of the time constraint, and ii) the distribution of escape durations for a fixed time constraint.

#### Adaptive stopping model

Since neither of these two features was explicitly represented in our model, we assumed that our ideal observer initiated a fast and inaccurate escape if its inference did not converge within the time limit. As above, we simulated the inference process for *K* = 100, 000 trials, and we took the fraction of trials that did not converge at or before *t* = *T*_max_. To estimate the “duration” of escape, we assumed that both early and late stops could be used to initiate an escape, and that the duration of this escape would last the remainder of the available inference time: i.e., duration = *T*_max_ − *t*_stop_, where *t*_stop_ is the time at which the inference converged.

#### Probabilistic speed-accuracy tradefoff (SAT) model

To construct a model for escape behavior based on the classic SAT, we assumed that the observer makes a decision at each time step *t* to either continue gathering data, made with probability *p*_sample_(*t*), or to stop the inference and initiate an escape, made with probability *p*_escape_(*t*) = 1 ™ *p*_sample_(*t*). We used the SAT curve (Fig. 2d) to specify the probability *p*_sample_(*t*); the larger the average error at time *t*, the higher the probability that the agent will continue to sample. Because the average error is roughly exponential as a function of time, we assumed that *p*_sample_(*t*) = exp(*λt*), where *λ* is a “difficulty parameter”. For *λ* = 0, the inference task is difficult, and inference always continues until the time limit *T*_max_. For *λ*→−∞, the inference task is easy, and the agent stops sampling immediately after the first sample is observed. For a given value of *λ*, one can analytically calculate the total fraction of converged inference trials for which the agent stopped sampling and initiated an escape within the time limit *T*_max_:

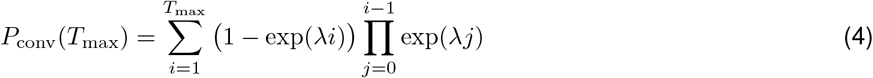

and conversely, the fraction of non-converged trials is given by:

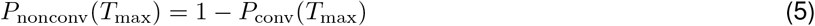

We used these calculations to plot the results in Fig. 6g.

## References

1. Stocker, A. A. & Simoncelli, E. P. Noise characteristics and prior expectations in human visual speed perception. Nature neuroscience 9, 578–585 (2006).

2. Młynarski, W. F. & Hermundstad, A. M. Adaptive coding for dynamic sensory inference. Elife 7, e32055 (2018).

3. Tavoni, G., Doi, T., Pizzica, C., Balasubramanian, V. & Gold, J. I. Human inference reflects a normative balance of complexity and accuracy. Nature human behaviour 6, 1153–1168 (2022).

4. Tavoni, G., Balasubramanian, V. & Gold, J. I. What is optimal in optimal inference? Current Opinion in Behavioral Sciences 29, 117–126 (2019).

5. Evans, D. A., Stempel, A. V., Vale, R. & Branco, T. Cognitive control of escape behaviour. Trends in cognitive sciences 23, 334–348 (2019).

6. Cooper, W. E. & Blumstein, D. T. Escaping from predators: an integrative view of escape decisions (Cambridge University Press, 2015).

7. Domenici, P., Blagburn, J. M. & Bacon, J. P. Animal escapology I: theoretical issues and emerging trends in escape trajectories. Journal of Experimental Biology 214, 2463–2473 (2011).

8. Card, G. & Dickinson, M. Performance trade-offs in the flight initiation of Drosophila. Journal of Experimental Biology 211, 341–353 (2008).

9. Heitz, R. P. The speed-accuracy tradeoff: history, physiology, methodology, and behavior. Frontiers in neuroscience 8, 86875 (2014).

10. Bogacz, R., Wagenmakers, E.-J., Forstmann, B. U. & Nieuwenhuis, S. The neural basis of the speed–accuracy tradeoff. Trends in neurosciences 33, 10–16 (2010).

11. Dambacher, M. & Hübner, R. Time pressure affects the efficiency of perceptual processing in decisions under conflict. Psychological research 79, 83–94 (2015).

12. Forstmann, B. U., Ratcliff, R. & Wagenmakers, E.-J. Sequential sampling models in cognitive neuroscience: Advantages, applications, and extensions. Annual review of psychology 67, 641–666 (2016).

13. Plamondon, R. & Alimi, A. M. Speed/accuracy trade-offs in target-directed movements. Behavioral and brain sciences 20, 279–303 (1997).

14. Drugowitsch, J., DeAngelis, G. C., Angelaki, D. E. & Pouget, A. Tuning the speed-accuracy trade-off to maximize reward rate in multisensory decision-making. elife 4, e06678 (2015).

15. Mendonça, A. G. et al. The impact of learning on perceptual decisions and its implication for speed-accuracy tradeoffs. Nature communications 11, 2757 (2020).

16. Rinberg, D., Koulakov, A. & Gelperin, A. Speed-accuracy tradeoff in olfaction. Neuron 51, 351–358 (2006).

17. Chittka, L., Dyer, A. G., Bock, F. & Dornhaus, A. Bees trade off foraging speed for accuracy. Nature 424, 388–388 (2003).

18. Latty, T. & Beekman, M. Speed–accuracy trade-offs during foraging decisions in the acellular slime mould Physarum poly-cephalum. Proceedings of the Royal Society B: Biological Sciences 278, 539–545 (2011).

19. Jaynes, E. T. Probability theory: The logic of science (Cambridge university press, 2003).

20. Card, G. M. Escape behaviors in insects. Current opinion in neurobiology 22, 180–186 (2012).

21. Simoncelli, E. P. Optimal estimation in sensory systems. The Cognitive Neurosciences, IV, 525–535 (2009).

22. Cover, T. M. Elements of information theory (John Wiley & Sons, 1999).

23. Touchette, H. A basic introduction to large deviations: Theory, applications, simulations. arXiv preprint 1106.4146 (2011).

24. Meyniel, F., Sigman, M. & Mainen, Z. F. Confidence as Bayesian probability: From neural origins to behavior. Neuron 88, 78–92 (2015).

25. Von Reyn, C. R. et al. Feature integration drives probabilistic behavior in the Drosophila escape response. Neuron 94, 1190–1204 (2017).

26. Von Reyn, C. R. et al. A spike-timing mechanism for action selection. Nature neuroscience 17, 962–970 (2014).

27. Eaton, R., Lee, R. & Foreman, M. The Mauthner cell and other identified neurons of the brainstem escape network of fish. Progress in neurobiology 63, 467–485 (2001).

28. Dai, Z., Yu, H., Low, B. K. H. & Jaillet, P. Bayesian optimization meets Bayesian optimal stopping in International conference on machine learning (2019), 1496–1506.

29. Arrow, K. J., Blackwell, D. & Girshick, M. A. Bayes and minimax solutions of sequential decision problems. Econometrica, Journal of the Econometric Society, 213–244 (1949).

30. Brockwell, A. E. & Kadane, J. B. A gridding method for Bayesian sequential decision problems. Journal of Computational and Graphical Statistics 12, 566–584 (2003).

31. Drugowitsch, J., Moreno-Bote, R., Churchland, A. K., Shadlen, M. N. & Pouget, A. The cost of accumulating evidence in perceptual decision making. Journal of Neuroscience 32, 3612–3628 (2012).

32. Evans, D. A. et al. A synaptic threshold mechanism for computing escape decisions. Nature 558, 590–594 (2018).

33. Młynarski, W. Efficient coding of spectrotemporal binaural sounds leads to emergence of the auditory space representation. Frontiers in computational neuroscience 8, 26 (2014).

34. Mugan, U. & MacIver, M. A. Spatial planning with long visual range benefits escape from visual predators in complex naturalistic environments. Nature communications 11, 3057 (2020).

35. Shamash, P. et al. Mice learn multi-step routes by memorizing subgoal locations. Nature neuroscience 24, 1270–1279 (2021).

36. Bitzer, S., Park, H., Blankenburg, F. & Kiebel, S. J. Perceptual decision making: drift-diffusion model is equivalent to a Bayesian model. Frontiers in human neuroscience 8, 102 (2014).

37. Tajima, S., Drugowitsch, J., Patel, N. & Pouget, A. Optimal policy for multi-alternative decisions. Nature neuroscience 22, 1503–1511 (2019).

38. Shadlen, M. N. & Newsome, W. T. Neural basis of a perceptual decision in the parietal cortex (area LIP) of the rhesus monkey. Journal of neurophysiology 86, 1916–1936 (2001).

39. Koblinger, Á., Fiser, J. & Lengyel, M. Representations of uncertainty: where art thou? Current Opinion in Behavioral Sciences 38, 150–162 (2021).

40. Temizer, I., Donovan, J. C., Baier, H. & Semmelhack, J. L. A visual pathway for looming-evoked escape in larval zebrafish. Current Biology 25, 1823–1834 (2015).

41. Murphy, K. P. Conjugate Bayesian analysis of the Gaussian distribution. arXiv (2007).

42. Evans, D. A., Stempel, A. V., Vale, R. & Branco, T. Cognitive control of escape behaviour. Trends in cognitive sciences 23, 334–348 (2019).

43. Domenici, P., Blagburn, J. M. & Bacon, J. P. Animal escapology II: escape trajectory case studies. Journal of Experimental biology 214, 2474–2494 (2011).

44. Gershman, S. J., Horvitz, E. J. & Tenenbaum, J. B. Computational rationality: A converging paradigm for intelligence in brains, minds, and machines. Science 349, 273–278 (2015).

45. Młynarski, W. F. & Hermundstad, A. M. Efficient and adaptive sensory codes. Nature Neuroscience 24, 998–1009 (2021).

